# Genetic correlations between pain phenotypes and depression and neuroticism

**DOI:** 10.1101/362574

**Authors:** Weihua Meng, Mark J Adams, Ian J Deary, Colin NA Palmer, Andrew M McIntosh, Blair H Smith

## Abstract

Correlations between pain phenotypes and psychiatric traits such as depression and the personality trait of neuroticism are not fully understood. The purpose of this study was to identify whether eight pain phenotypes, depressive symptoms, major depressive disorders, and neuroticism are correlated for genetic reasons. Eight pain phenotypes were defined by a specific pain-related question in the UK Biobank questionnaire. First we generated genome-wide association summary statistics on each pain phenotype, and estimated the common SNP-based heritability of each trait using GCTA. We then estimated the genetic correlation of each pain phenotype with depressive symptoms, major depressive disorders and neuroticism using the the cross-trait linkage disequilibrium score regression (LDSC) method integrated in the LD Hub. Third, we used the LDSC software to calculate genetic correlations among pain phenotypes. All pain phenotypes were heritable, with pain all over the body showing the highest heritability (*h^2^*=0.31, standard error=0.072). All pain phenotypes, except hip pain and knee pain, had significant and positive genetic correlations with depressive symptoms, major depressive disorders and neuroticism. The largest genetic correlations occurred between neuroticism and stomach or abdominal pain (rg=0.70, *P*=2.4 x 10^−9^). In contrast, hip pain and knee pain showed weaker evidence of shared genetic architecture with these negative emotional traits. In addition, many pain phenotypes had positive and significant genetic correlations with each other indicating shared genetic mechanisms. Pain at a variety of body sites is heritable and genetically correlated with depression and neuroticism. This suggests that pain, neuroticism and depression share partially overlapping genetic risk factors.

## INTRODUCTION

Pain is a global public health priority. In the Global Burden of Diseases Study 2015, low back and neck pain were, by far, the leading cause of years lived with disability (YLDs) worldwide, with migraine also ranked among the top ten causes.^1^ Some other leading causes of YLDs such as musculoskeletal disorders and diabetes are also highly likely to feature pain as a prominent symptom. It was estimated that 20% of adults suffer from pain globally and that 10% adults are diagnosed with chronic pain each year.^2^ Chronic pain, i.e. pain that has persisted beyond normal tissue healing time (usually taken as 3 months) can arise from many causes, but is often idiopathic or difficult to classify pathophysiologically.^3^ It is recognized to have significant genetic contributions to its development,^4^ as well as environmental components. Improving our understanding of the genetic contributions to the experience of pain could help our understanding of its aetiology and prevention, and might also offer opportunities for personalised medicine and drug targeting based on identified genetic variants. Although some genetic studies have focused on pain in specific body sites and have suggested possible genetic variants associated with pain phenotypes,^5^ the overall understanding of the genetic contributions of pain remain unclear. Current limitations in our knowledge include: 1. the extent to which pain as a phenotype is determined by additive genetic components mainly represented by single nucleotide polymorphisms (SNPs); 2. whether the genetic mechanisms of pain in different body sites or in different disorders are similar or different; and 3. whether the genetic connections between pain phenotypes and other common co-morbidities are similar or different. Addressing these questions brings further challenges when the severity and the frequency of pain are taken into account.

Mental health disorders are among the commonest co-morbidities of pain.^6-7^Depression was ranked as the third most important cause of disability worldwide, and anxiety is also a leading cause.^1^ The personality trait of neuroticism shows a high phenotypic and genetic association with depression, anxiety and with reports of physical and mental disorders.^8-10^ Neuroticism is also reckoned to cause a large burden of ill health and health-related expenditure.^11^ Specific pain phenotypes including migraine, facial pain, neck pain, abdominal pain, low back pain, and fibromyalgia have been shown to be associated with depression and anxiety.^12-17^This epidemiological coexistence between pain and psychiatric traits could arise in part because of shared genetic factors.^18-19^ Understanding the genetic correlation between pain and psychiatric traits may help to elucidate their degree of shared genetic architecture and provide a framework for future causal inference.^20^ It has been proposed that some pain phenotypes (such as migraine, back pain) and some psychiatric traits (such as depression or neuroticism) share common genetic components.^21-24^ However, the genetic correlations between multiple pain phenotypes in different body sites, and those between pain phenotypes and depressive symptoms, major depressive disorders and neuroticism, have not been reported systematically, to the best of our knowledge.

The recently released UK Biobank cohort has recorded eight broadly-defined pain phenotypes, generated from a specific pain question: headache, facial pain, neck or shoulder pain, back pain, stomach or abdominal pain, hip pain, knee pain, and pain all over the body.^25^ This has made it possible for researchers to identify genetic variants associated with each pain phenotype. In addition, when combining genome-wide association study (GWAS) summary statistics with the cross-trait linkage disequilibrium score regression (LDSC) method implemented in the online LD hub tool (http://ldsc.broadinstitute.org/ldhub/) and the LDSC software (https://github.com/bulik/ldsc), researchers can estimate the genetic correlations between selected phenotypes.^26^

In order to identify genetic correlations between pain phenotypes and depression and the personality trait of neuroticism, as well as the genetic correlations among pain phenotypes, we generated GWAS summary statistics on eight pain phenotypes in different body sites based on the UK Biobank cohort and adapted the cross-trait LDSC method through the LD hub and the LDSC software.

## MATERIALS AND METHODS

### Participants

Over 500,000 people aged between 40 and 69 years were recruited by the UK Biobank cohort in 2006-2010 across England, Scotland and Wales. A detailed clinical, demographic, and lifestyle questionnaire was completed by all participants. Biological samples (blood, urine and saliva) were also provided for future analysis. Further information on the UK Biobank cohort can be found at www.ukbiobank.ac.uk. Ethical approval was granted by the National Health Service National Research Ethics Service (reference 11/NW/0382). The current study was conducted under approved UK Biobank data application number 4844.

DNA extraction and quality control (QC) were standardized and the detailed method can be found at http://www.ukbiobank.ac.uk/wp-content/uploads/2014/04/DNA-Extraction-at-UK-Biobank-October-2014.pdf. Genotyping was obtained from the bespoke Affymetrix UK Biobank chips. The Wellcome Trust Centre for Human Genetics at Oxford University was in charge of standard QC procedures for genotyping results. The detailed QC steps can be found at http://biobank.ctsu.ox.ac.uk/crystal/refer.cgi?id=155580.

In March 2018, The UK Biobank released an updated version of the genetic information (including directly genotyped genotypes and imputed genotypes) of 501,708 samples to all approved researchers. The detailed QC steps of imputation were described by Bycroft et al.^27^

### Definitions of pain phenotypes

We used a specific pain-related questionnaire adapted by the UK Biobank, which included the question: ‘In the last month have you experienced any of the following that interfered with your usual activities?’. The options were: 1. Headache; 2. Facial pain; 3. Neck or shoulder pain; 4. Back pain; 5. Stomach or abdominal pain; 6. Hip pain; 7. Knee pain; 8. Pain all over the body; 9. None of the above; 10. Prefer not to say. More than one option could be selected. (UK Biobank Questionnaire field ID: 6159)

For each pain phenotype, cases were defined as those who selected the specific pain site option for the above question, regardless of whether they had selected other options. For example, headache cases are those who selected the ‘Headache’ option; Facial pain cases are those who selected the ‘Facial pain’ option; etc. For each GWAS analysis, controls were those who selected the ‘None of the above’ option. Thus we used the same ‘no pain’ control population for all pain phenotypes in different body sites.

### Definitions of depression and neuroticism

The phenotypes of depression and neuroticism were defined by the psychological cohorts collected by the LD hub.^28-31^ The original researchers of these cohorts agreed to share the GWAS summary statistics on depression and neuroticism with the LD hub for generating genetic correlations. Therefore, we selected the ‘Psychiatric diseases’ option and the ‘Personality traits’ option in the LD hub to include the depression and neuroticism traits. These traits are: Depressive symptoms,^28^ Neuroticism (x 2 studies),^29-30^ Major depressive disorder.^31^ However, for neuroticism, we only chose the version used by Okbay et al,^30^ as it is a GWAS meta-analysis publication, the results of which included the results from van den Berg et al.^29^

### Statistical analysis

#### Generating the heritabilities of all pain phenotypes

In this study, genome-wide complex trait analysis (GCTA) was used to calculate narrow-sense SNP-based heritabilities based on the genomic-relatedness-based restricted maximum-likelihood (GREML) approach.^32^

#### Generating GWAS summary statistics of all pain phenotypes

In this study, genotype data were analysed in BGENIE (https://jmarchini.org/bgenie/), as recommended by UK Biobank. Routine QC steps included: removing SNPs with INFO scores less than 0.1, SNPs with minor allele frequency less than 0.5%, or SNPs that failed Hardy-Weinberg tests *P* < 10^−6^. SNPs on the X and Y chromosomes and mitochondrial SNPs were also removed. We further removed those whose ancestry was not white British based on principal component analysis, those who were related at least another participant in the cohort (a cut-off value of 0.025 in the generation of the genetic relationship matrix) and those who failed QC. Association tests based on standard Frequentist association were performed using BGENIE adjusting for age, sex, body mass index (BMI), 9 population principal components, genotyping arrays, and assessment centers.

#### Generating genetic correlations between pain phenotypes and depression and neuroticism by the LD hub

The LD hub has gathered 235 published GWAS summary statistics of different disorders worldwide. Those GWAS summary statistics were compared against researcher-uploaded GWAS summary statistics of a phenotype of interest to generate genetic correlations between the phenotype and 235 phenotypes.^26^

In order to identify genetic correlations between pain phenotypes and the depression and neuroticism traits, we used the cross-trait LDSC method through the LD Hub v1.9.0.^26^ The LD Hub estimates the bivariate genetic correlations of a phenotype with other traits using individual SNP allele effect sizes and the average LD in a region. In this study, as there are altogether three depression and neuroticism traits available in the LD hub, those with *P* values less than 0.002 (0.05/24, 8 pain phenotypes and 3 psychiatric phenotypes) should be considered significant surviving Bonferroni correction for multiple testing.

#### Generating genetic correlations among pain phenotypes using the LDSC software

The LDSC software is a Linux version of the online LD hub tool. It is used to calculate the genetic correlations between two traits at a time while the LD hub can output the genetic correlations between one trait and many traits at the same time. In this study, as there were altogether 28 pair combinations among eight pain phenotypes, those with *P* values less than 0.0018 (0.05/28) should be considered significant surviving Bonferroni correction for multiple testing.

## RESULTS

### The heritabilities and the GWAS summary statistics of all pain phenotypes

The specific pain question received 775,252 responses to all options answered by 501,708 UK Biobank participants during the initial assessment visit (2006-2010). Table 1 summarises the numbers of cases and controls in the GWAS of the eight pain phenotypes.

**Table 1.**
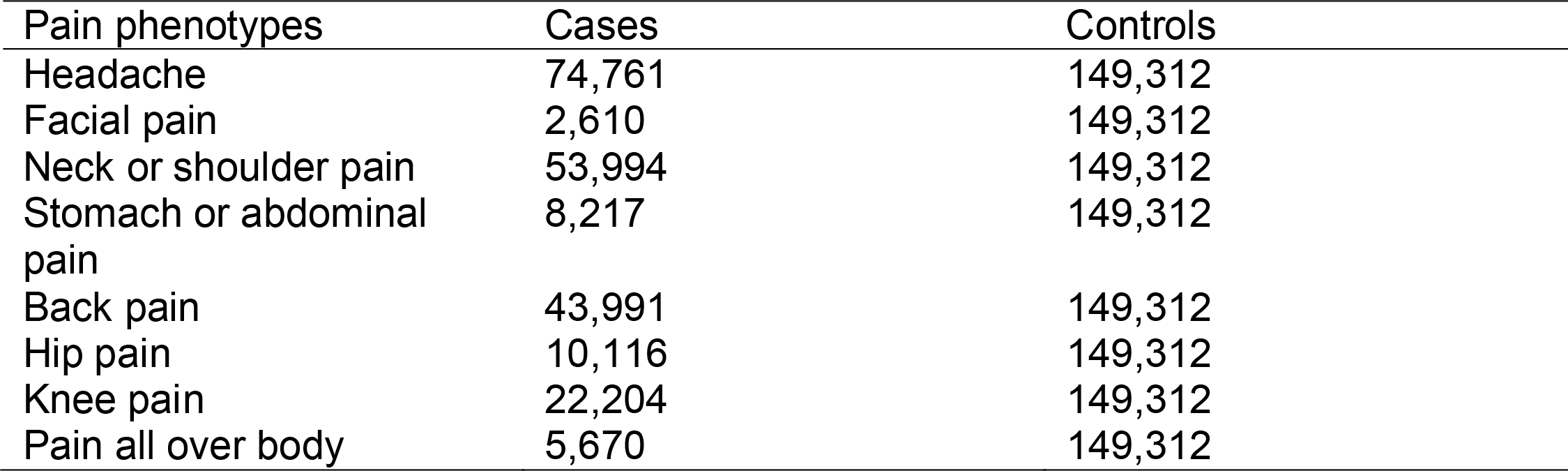
The sample numbers available for GWAS on eight pain phenotypes.

The narrow-sense SNP heritabilities of each pain phenotype are presented in the Table 2. Pain all over the body had the greatest heritability among all pain phenotypes (*h^2^*=0.31, standard error (SE)=0.072). Knee pain has the lowest heritability (*h^2^*=0.08, SE=0.029). The SNP heritabilities of other pain phenotypes were between 0.11 to 0.24.

**Table 2.** The SNP-based heritabilities (*h^2^*) of eight pain phenotypes from the UK Biobank cohort and their genetic correlations with depressive symptoms, major depression and neuroticism.

| Pain phenotypes | Heritability |  | Depressive symptoms |  | Major depressive disorder |  | Neuroticism |  |
| --- | --- | --- | --- | --- | --- | --- | --- | --- |
| | $h^2$ (SE) | $P$ | $rg$ | $P$ | $rg$ | $P$ | $rg$ | $P$ |
| Headache | 0.21<br>(0.015) | $1.6 \times 10^{-206}$ | **0.52** | $1.6 \times 10^{-46}$ | **0.39** | $1.6 \times 10^{-11}$ | **0.50** | $2.2 \times 10^{-72}$ |
| Facial pain | 0.24<br>(0.12) | $3.2 \times 10^{-41}$ | **0.33** | $2 \times 10^{-4}$ | 0.34 | 0.01 | **0.30** | $1.0 \times 10^{-5}$ |
| Neck or shoulder pain | 0.11<br>(0.017) | $1.3 \times 10^{-153}$ | **0.55** | $3.4 \times 10^{-30}$ | **0.40** | $5.8 \times 10^{-8}$ | **0.44** | $5.3 \times 10^{-7}$ |
| Stomach or abdominal pain | 0.14<br>(0.050) | $6.5 \times 10^{-54}$ | **0.67** | $5.7 \times 10^{-7}$ | **0.53** | $5 \times 10^{-4}$ | **0.70** | $2.4 \times 10^{-9}$ |
| Back pain | 0.11<br>(0.020) | $1.5 \times 10^{-67}$ | **0.48** | $1.5 \times 10^{-14}$ | **0.36** | $3 \times 10^{-5}$ | **0.40** | $1.7 \times 10^{-13}$ |
| Hip pain | 0.12<br>(0.041) | $4.5 \times 10^{-56}$ | 0.34 | 0.03 | 0.04 | 0.80 | 0.27 | 0.04 |
| Knee pain | 0.08<br>(0.029) | $4.1 \times 10^{-56}$ | 0.12 | 0.13 | -0.07 | 0.53 | 0.18 | 0.002 |
| Pain all over body | 0.31<br>(0.072) | $5.9 \times 10^{-74}$ | **0.69** | $1.4 \times 10^{-27}$ | **0.43** | $5.6 \times 10^{-6}$ | **0.45** | $3.4 \times 10^{-17}$ |

$P$ values $< 0.002$ (0.05/24) were considered as significant. Those significant $rg$ values were in bold.
SE: standard error
$rg$ : genetic correlation

### Genetic correlations between pain and depression and neuroticism

Through the genetic correlation analysis, we identified multiple significant and positive correlations between pain phenotypes and depression and neuroticism. (Table 2, Figure 1) A supplementary table is provided to show the genetic correlations between pain phenotypes and all available psychiatric and personality traits in the LD hub (Supplementary Table 1).

**Figure 1.**
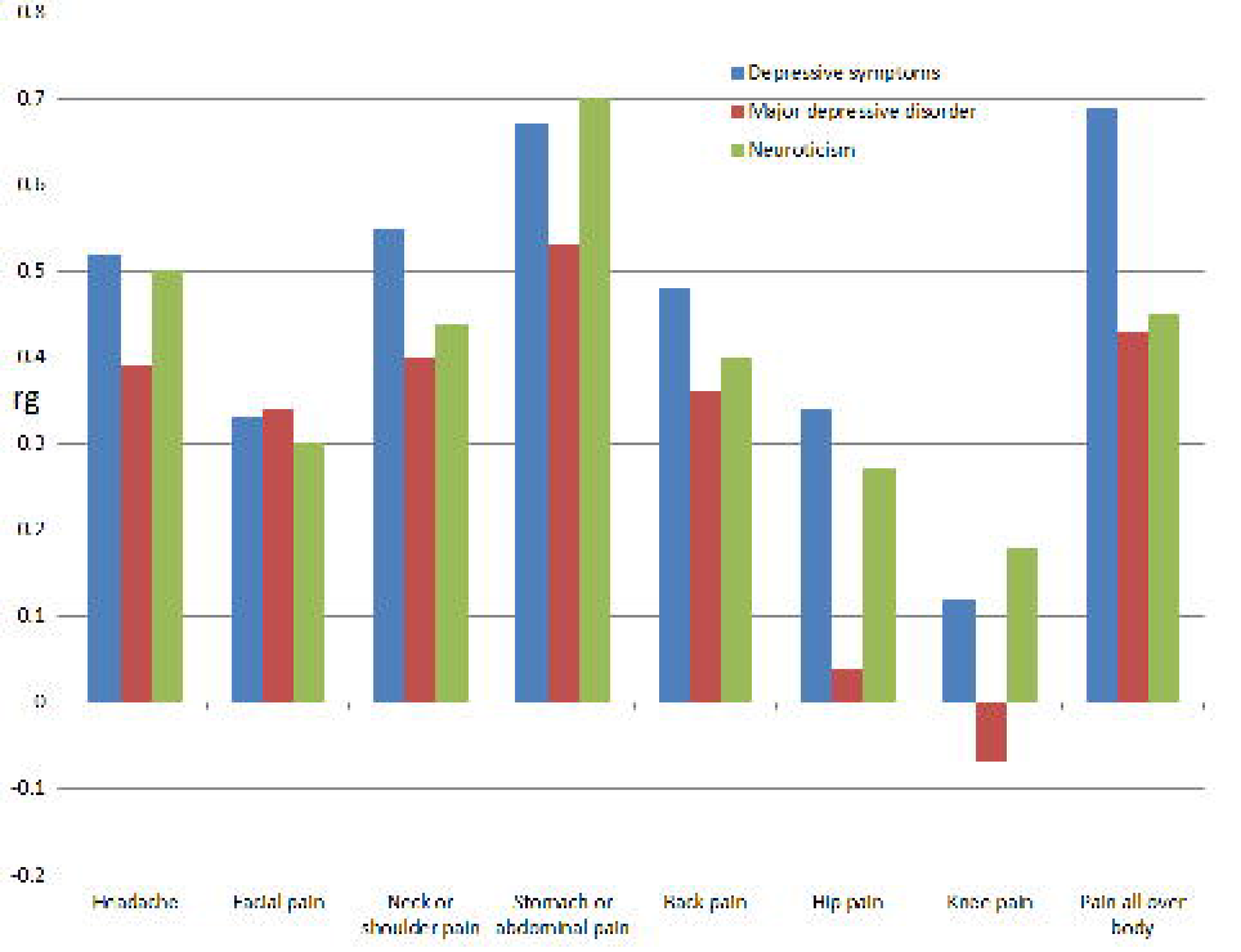
The genetic correlations between eight pain phenotypes and depressive symptoms, major depressive disorders and neuroticism Please note, the genetic correlations between these traits and hip pain and knee pain were not significant (*P* > 0.002, Table 2). Rg: genetic correlation

#### Depression and eight pain phenotypes

For depressive symptoms, all pain phenotypes had significant and positive genetic correlations with depression except hip pan and knee pain. The largest genetic correlation occurred with pain all over the body (rg=0.69, *P*=1.4 x 10^−27^), followed by stomach or abdominal pain (rg=0.67, *P*=5.7 x 10^−7^). For hip pain and knee pain, although there were positive genetic correlations with depressive symptoms (rg=0.34 and 0.12, correspondingly), the associations did not survive Bonferroni correction (*P*=0.03 and 0.13, correspondingly). The values of the genetic correlations between other pain phenotypes and depressive symptoms were between rg=0.33 and 0.55, and all were statistically significant after adjustment for multiple testing.

For major depressive disorder, the genetic correlation results were similar to those for depressive symptoms. The largest genetic correlation was with stomach or abdominal pain (rg=0.53, *P*=0.0005), followed by pain all over the body (rg=0.43, *P*=5.6 x 10^−6^). However, the rg values of hip pain and knee pain were 0.04 and −0.07, which were also statistically insignificant (*P*=0.80 and 0.53, correspondingly). The values of the genetic correlations between other pain phenotypes and major depressive disorder were between rg=0.34 and 0.40, and all were statistically significant.

#### Neuroticism and eight pain phenotypes

With neuroticism, stomach or abdominal pain had the largest genetic correlation (rg=0.70, *P*=2.4 x 10^−9^). Headache followed next with rg=0.50 and *P*=2.2 x 10^−72^. All genetic correlations with other pain phenotypes (except hip pain and knee pain) were positive and significant with rg values between 0.30 and 0.50. For hip pain and knee pain, although there were positive genetic correlations with neuroticism, the correlations were statistically insignificant.

### Genetic correlations among pain phenotypes

Through the LDSC software, we identified multiple significant and positive correlations among pain phenotypes (Table 3). The largest positive and significant genetic correlation was between neck or should pain and back pain (rg=0.83, *P*=2.11 x 10^−100^), followed by hip pain and pain all over the body (rg=0.81, *P*=0.0004). Neck or shoulder pain had positive and significant genetic correlations with all other pain phenotypes (0.52<rg<0.83), this was the same for pain all over the body(0.36<rg<0.81). Among all all pain phenotypes, hip pain only had three positive and significant genetic correlations (neck or shoulder pain, back pain and pain all over the body) with other pain phenotypes.

**Table 3.** The genetic correlations among all pain phenotypes based on the UK Biobank.

|  | Headache |  | Facial pain |  | Neck or shoulder pain |  | Stomach or abdominal pain |  | Back pain |  | Hip pain |  | Knee pain |  |
| --- | --- | --- | --- | --- | --- | --- | --- | --- | --- | --- | --- | --- | --- | --- |
|  | rg | P | rg | P | rg | P | rg | P | rg | P | rg | P | rg | P |
| Facial pain | **0.44** | 1.06 x 10<sup>-10</sup> |  |  |  |  |  |  |  |  |  |  |  |  |
| Neck or shoulder pain | **0.61** | 1.39 x 10<sup>-94</sup> | **0.52** | 3.27 x 10<sup>-10</sup> |  |  |  |  |  |  |  |  |  |  |
| Stomach or abdominal pain | **0.56** | 8.01 x 10<sup>-10</sup> | **0.60** | 9.16 x 10<sup>-5</sup> | **0.67** | 3.3 x 10<sup>-12</sup> |  |  |  |  |  |  |  |  |
| Back pain | **0.53** | 2.79 x 10<sup>-36</sup> | **0.47** | 4.50 x 10<sup>-7</sup> | **0.83** | 2.11 x 10<sup>-100</sup> | **0.56** | 5.61 x 10<sup>-6</sup> |  |  |  |  |  |  |
| Hip pain | 0.32 | 0.002 | 0.86 | 0.002 | **0.69** | 1.79 x 10<sup>-5</sup> | 0.38 | 0.12 | **0.54** | 0.0004 |  |  |  |  |
| Knee pain | **0.39** | 3.70 x 10<sup>-13</sup> | 0.33 | 0.01 | **0.53** | 1.53 x 10<sup>-14</sup> | **0.53** | 0.0002 | **0.46** | 3.29 x 10<sup>-8</sup> | 0.36 | 0.03 |  |  |
| Pain all over body | **0.62** | 4.69 x 10<sup>-52</sup> | **0.43** | 6.51 x 10<sup>-5</sup> | **0.79** | 1.45 x 10<sup>-47</sup> | **0.73** | 2.58 x 10<sup>-7</sup> | **0.69** | 1.46 x 10<sup>-29</sup> | **0.81** | 0.0004 | **0.36** | 5.56 x 10<sup>-5</sup> |

P values < 0.0018 (0.05/28) were considered as significant. Those significant rg values were in bold.
rg: genetic correlation

## DISCUSSION

Eight self-reported pain phenotypes, from different sites across the body, were all heritable and showed a broad pattern of partially shared genetic architecture with each other. We also found evidence that pain shared genetic architecture with depressive symptoms, major depressive disorder and neuroticism for most sites across the body. Hip pain and knee pain was the exception, in showing weak and non-significant genetic correlations with depressive symptoms, major depressive disorder and neuroticism.

The nature of the relationship between pain phenotypes and depression has been uncertain. Epidemiological studies have identified that depression is reported more often by patients reporting pain and also that pain is a risk factor for the future development of depression.^33^ Patients with psychosomatic conditions suffer from psychiatric disorders (such as depression) and physical diseases (such as bodily pain), and their co-occurrence is associated with greater disability than either condition alone.^34^ Although the exact mechanisms linking these conditions are not clear, genetic mechanisms are implied through shared biological pathways, such as gene expression in biological networks, the endocannabinoid system, the hypothalamic-pituitary-adrenal axis and inflammatory pathways.^35^ Our study answered a specific question about depression: to what extent is each pain phenotype genetically correlated with depression? Our genetic correlation results between headache, facial pain, neck or shoulder pain, stomach or abdominal pain, back pain, pain all over the body and depression are all consistent in direction with known epidemiological associations.^12-17^ This suggests that share genetic risk factors are likely to partly explain their phenotypic correlations. However, the genetic correlations between hip pain and knee pain and depression were contrary to previous observations that depression and knee pain or hip pain are strongly related.^36^ Previous studies have shown that knee pain from osteoarthritis increases a person’s risk of developing subsequent depression.^37^ A systematic review of the relationship between knee pain and multiple psychiatric traits also found an association between depression and knee pain.^38^ While genetic factors may contribute to the pain at different sites, our findings suggest that non-genetic factors may be more important in the co-occurrence of knee or hip pain with depression. The genetic relationships between pain phenotypes and neuroticism are also of interest and are similar to those between pain and depression. Neuroticism was identified to be a potential risk factor for elevated pain responses in laboratory pain in healthy children, and can likely exacerbate pain responses when coupled with fear of bodily sensations.^39^ Neuroticism has also been independently associated with greater pain catastrophizing and pain-related anxiety.^40^ Our genetic correlation results were consistent with findings from epidemiological studies of headache,^41^neck or shoulder pain,^42^ back pain,^43^ and pain all over the body.^44^ No previous studies have shown epidemiological data for the relationships between neuroticism and facial pain or stomach or abdominal pain. It was also suggested from our study that there were no significant genetic relationships between knee pain or hip pain and neuroticism. To our knowledge, no previous studies have specifically examined the epidemiological relationships between neuroticism and hip pain or knee pain. Our findings suggest that hip pain and knee pain may belong to a separate pain group and should be considered separately when designing studies of the genetic relationships between pain and psychiatric disorders.

A significant and positive correlation between a pain phenotype and a psychiatric trait reflects share genetic architecture. This can reflect pleiotropy, where specific genetic alleles give risk to both phenotypes, but it can also reflect mediated pleiotropy where there is a directional or causal association between these traits.^20^ It is also possible for positive genetic correlations to be generated by misclassification of pain as depression, or major depression as pain.^45^ Whilst this seems unlikely to explain the convergent genetic correlation of neuroticism with pain, it will be important to examine whether there is subgroup heterogeneity amongst individuals with pain on depression that can be explained by variation on the other trait.

This paper also reported the genetic correlations among pain phenotypes in different body sites. Many pain phenotypes have positive and significant genetic correlations with each other indicating the common genetic mechanisms behind different pain phenotypes. This common mechanism is less likely to be represented by a few genes with large effects, but to reflect many genetic variants with smaller effects. It is biologically plausible for back pain and neck or shoulder pain to demonstrate the largest genetic correlation (rg=0.83) since causal genetic factors could have plausible detrimental effects across the whole spine. For other genetic correlations such as that between hip pain and pain all over the body (rg=0.81), the reason is less apparent and merits further research.

The highest narrow-sense heritability among all pain phenotypes in this study was 0.31 for pain all over the body. The heritablities of all other pain phenotypes were moderate. The narrow-sense heritability does not take gene-gene interactions, gene-environment interactions, or the contribution from rare variants into account, and is therefore likely to be an under estimate of the true heritability. This is the first report of the heritabilities for facial pain (*h^2^*=0.24), stomach or abdominal pain (*h^2^*=0.14), to the best of our knowledge, and suggests important genetic contributions to chronic pain at all body sites.

Arguably the greatest strength of the current study was the large size of the UK Biobank sample. This provided the largest single sample size for many of the pain phenotypes studied here compared with previous GWAS studies of pain.^5^ Nevertheless, potential limitations should also be taken into account. The phenotyping in UK Biobank was based on a single specific non-standard pain-related question. This means that all pain phenotypes were broadly-defined and unfiltered by other potentially relevant information on the nature, duration or intensity of the pain. Similar limitations also apply to the psychological traits measured in UK Biobank and elsewhere.^46^ In future, new pain questionnaires may be administered to participants in UK Biobank, and this will allow for more detailed and focused phenotyping for use in future analyses.

In summary, we have identified significant and positive genetic correlations between multiple pain phenotypes and depression and neuroticism, suggesting that the known associations between these traits are partly due to shared genetic architecture. In contrast, we have suggested that the known epidemiological relationships between hip and knee pain and depression are not caused primarily by common genetic factors, prompting a search for other explanations. In addition, we have showed that many pain phenotypes are heritable and have positive and significant genetic correlations with each other. This indicates that common genetic risk factors confer liability to pain at many different sites across the body, suggesting shared risk factors and, potentially, disease mechanisms.

These findings contribute to the understanding of the genetic and biological mechanisms for individual pain phenotypes, depression and neuroticism. In addition, the findings also represent an early but important step towards the identification of causal associations between pain phenotypes and psychiatric disorders and identifying subgroup heterogeneity.

## ACKNOWLEDGEMENTS

We would like to thank all participants of the UK Biobank cohort who have provided necessary genetic and phenotypic information.

This work was supported by the STRADL project [Wellcome Trust, grant number: 104036/Z/14/Z], and the Centre for Cognitive Ageing and Cognitive Epidemiology [Medical Research Council and Biotechnology and Biological Sciences Research Council, grant number: MR/K026992/1]. We are grateful for support from the Sackler Foundation. The funders had no role in study design, data collection, data analysis, interpretation, writing of the report.

